# Thermal Pain and Detection Threshold Modulation in Augmented Reality

**DOI:** 10.1101/2022.06.22.497278

**Authors:** Daniel Eckhoff, Christian Sandor, Gladys L. Y. Cheing, Jan Schnupp, Alvaro Cassinelli

## Abstract

Augmented Reality (AR) overlays computer-generated visual, auditory or other sensory information on the real world. Due to recent advancements in AR it can be difficult for the user to differentiate between sensory information coming from real and virtual objects, leading to interesting phenomena. For example, an AR experience in which users can experience their own hands in flames has been shown to elicit heat illusions on the affected hands. In this study, we investigate the potential that AR has for top-down modulation of pain and thermal perception. We assessed thermal pain and detection thresholds on the participant’s right hand when they experienced that hand covered in virtual flames in AR. We compared that experience to a baseline condition with no additional stimuli as well to a control condition that sees the hand covered by unrealistic blue flames to compensate the distraction factor. We found that experiencing a burning hand in AR induced analgesic and hyperalgesic effects as participants began to feel heat related pain on lower temperatures and cold related pain on higher temperatures. That experience also significantly changed the lowest temperature at which participants started perceiving warmth. These results demonstrate that pain and thermal perception can be manipulated by altering the perception of our body in AR.

## 1 INTRODUCTION

Augmented Reality (AR) is an emerging interactive technology that changes the user’s perception of the real world by adding virtual sensory stimuli on the real world, or through the modification or removal of sensory information from real objects. When these virtual sensory stimuli are coherent to the real world and experienced in an immersive manner (e.g, through head-worn displays), they can be indistinguishable from actual real sensory information, and can therefore prompt the natural associated response. For example, Weir and colleagues have reported that an AR experience during which users experience their own hands covered in flames create involuntary heat and olfactory illusions (Weir et al., 2013). This raises the question of the extent to which this AR experience can modulate thermal perception. In this work we set to investigate to what extent AR modulate both thermal pain and detection thresholds in healthy volunteers. In particular, we want to address the following research questions: **RQ 1:** Does displaying virtual flames on the user’s hand lead to a change in Cold Pain Thresholds (CPT) and Heat Pain Thresholds (HPT)? **RQ 2:** Does displaying virtual flames on the user’s hand lead to a change in Cold Detection Thresholds (CDT) or Warm Detection Thresholds (WDT)?

The major finding of the present study was that AR does indeed modulate nociception and perception of real noxious and non-noxious thermal stimuli. Experiencing the virtual burning hand significantly **(1)** lowers HPT and CPT as well as **(2)** WDT, but not CDT. Our control condition did not change thresholds significantly compared to baseline, suggesting that the changes in thresholds do not come from possible distracting factors.

Thermal perception thresholds are the temperatures at which a person begins to perceive a thermal stimulus as warm or cold, while thermal pain thresholds are the temperatures at which a person begins to perceive a heat or cold stimulus as painful. These thresholds are usually determined by applying a thermal stimulator to the skin, slowly increasing or decreasing the temperature until the subject signals a thermal sensation or a pain induced by heat or cold respectively. Threshold values can depend on a number of factors, including the method of testing (Defrin et al., 2006), the rate of heating, (Yarnitsky and Ochoa, 1990) the ambient temperature (Strigo and Carli, 2000), the location on the skin (Defrin et al., 2006) or skin temperature (Pertovaara et al., 1996). Thresholds can also vary as a function of the participant’s, gender (Averbeck et al., 2017), or medical condition (Curković et al., 1993).

Cognitive factors or cross-modal interactions can also greatly modulate the perception of pain. (Longo et al., 2009) reported that the vision of one’s own body part in pain reduces the pain intensity. They delivered potentially painful stimuli to a participant’s hand while the participants were looking at their own hand, a box, or somebody else’s hand. Looking at one’s own had was associated with the lowest reported pain ratings.

Similar results have been shown in the Rubber Hand Illusion (RHI) (Botvinick and Cohen, 1998), where participant’s are embodying a rubber hand as their own. (Valenzuela-Moguillansky et al., 2011) showed how pain ratings of heat stimuli decreased immediately after the RHI.

Virtual Reality (VR) has been the object of a lot of research in pain perception. In contrast to AR, VR lets the user perceive only virtual sensory information, without information from the real world. In the near future, virtual stimuli presented in AR or VR may be barely distinguishable from real ones. Already, AR and VR experiences can induce significant feelings of presence, making people respond as if the experience was real (Slater, 2009; Sanchez-Vives and Slater, 2005). The sense of ‘being there’ - immersed in a virtual environment - is responsible for a significant level of distraction, which is why VR has attracted much research on pain perception. VR has been shown to be able to produce strong analgesic effects. For example, Hoffman placed patients suffering from burn injuries in a virtual snow world and demonstrated analgesic effects through subjective and fMRI measurements (Hoffman et al., 2000, 2004). However, the importance of the type of virtual environment is unclear. For example, (Mühlberger et al., 2007) had participants move through a snowy virtual winter landscape as well as through a predominantly yellow and red autumn landscape, and found similar analgesic effects in both environments, hinting at the possibility that analgesic effects may mostly rely on distracting the participant’s attention away from their pain. A few articles examined the role of distraction in VR on pain perception by manipulating the degree of cognitive load. Surprisingly, (Demeter et al., 2018) reported that there was no significant difference in pain reduction between VR experiences with high and low cognitive load. Both reduced the pain perception compared to a control condition. A subsequent study by (Barcatta et al., 2022) found that participants who reported more emotional distress exhibited higher pain thresholds with lower cognitive load. They highlight that individual differences should be considered when designing VR treatments for analgesic purposes.

Other studies have examined how pain perception is modulated by the appearance and interaction of embodied body parts, i.e., a virtual representation of a limb, collocated with the real limb in egocentric space, in VR. (Martini et al., 2013) used a VR system that superimposed a red, blue, or green color on a virtual embodied arm in VR. In their experiment, the reddened skin of the arm slightly decreased HPT, while the blue skin increased it. The experimental conditions hint at a different mechanism for pain modulation where distraction plays a secondary role compared to virtual embodiment. The effect of embodiment in VR on pain perception has been the subject of a number of studies (Hoffman, 2021; Käthner et al., 2019; Martini et al., 2015; Martini, 2016).

(Gandy et al., 2010) pointed out a crucial difference between AR and VR systems: the ability of the participant to observe their own body and its movement in real-time, which is not possible in VR. In VR, the experience can be either fully disembodied, or participants can embody limbs of a virtual avatar. However, virtual embodiment is a fragile perceptual experience not experienced by every subject to a full degree (Bekrater-Bodmann et al., 2012; Kilteni et al., 2012). Furthermore, it is difficult to integrate VR into everyday life because the real world cannot be perceived at the same time. So far, AR did mostly receive attention as a pain treatment for phantom limb pain (Carrino et al., 2014; Dunn et al., 2017; Ortiz-Catalan et al., 2016). Even though AR allows possible pain interventions to be seamlessly integrated into the daily life of patients.

## 2 MATERIALS AND METHODS

To investigate the effect of a visual-auditory illusion of fire on cutaneous heat sensation, we used thermal quantitative sensory tests (QSTs) to measure temperature thresholds for noxious and non-noxious thermal stimuli applied to the skin concurrently with three different visual-auditory conditions: realistic fire, unrealistic fire, and a control condition without AR manipulation of visual-auditory input. This allowed us to examine quantitatively how thermal sensitivity and pain thresholds were modulated by our AR system.

We designed a within-subject study that comprised two different visual and auditory conditions, as well as a baseline condition without additional stimuli. All conditions involved participants wearing a head-mounted AR display. We asked all 21 participants to keep looking at their dominant right hand while we measured the four different QST outcomes on that hand: WDT, CDT, HPT, and CPT. The setup and stimulus conditions are illustrated in Figure 1. In the “realistic fire” condition, participants experienced their own hands burning (Figure 1**(c)**). This involved a fire simulation that tracked the shape and movement of the participant’s hand, and which was complemented with sound effects of a burning torch. We also introduced another “unrealistic fire” control condition in order to be able to investigate whether highly realistic looking flames covering the hand were necessary to elicit any observed threshold changes, or whether other AR illusions with similar dynamics would be just as effective (Figure 1**(d)**). This control condition closely resembled the realistic fire in many respects, engulfing the hand in an identical way and moving with essentially identical dynamics, but the control condition flames moved in an unrealistic direction (i.e. downwards) and had a blue color that is atypical for flames emanating from burning materials which are rich in carbon. These control visuals were complemented with sound effects of howling winds, rather than those of a burning torch. We compared these two conditions to a baseline condition in which participants observed their right hand through the AR headset, but without any superimposed visual and auditory stimuli.

**Figure 1.**
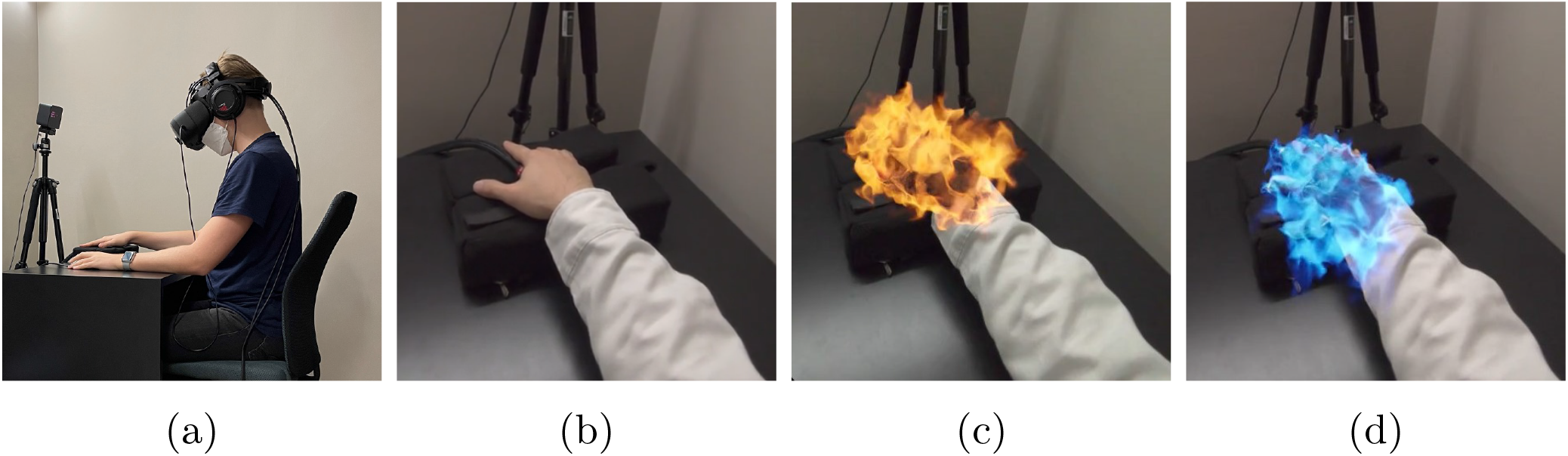
The experimental set-up and the three experimental conditions: the participant sees the graphics covering their hand through the AR headset. **(a)** Side view of the the participant wearing the AR headset, left arm over a computer mouse, right arm resting on a cushion, with the thermal stimulator in contact with the thenar prominence of the right hand. **(b)** Baseline condition. Participant is looking through the headset on their right hand with no additional auditory and visual stimuli. **(c)** Experiencing their own hand burning. **(d)** Experiencing blue flame like visuals covering their hand.

### 2.1 Participants

We recruited twenty-four right-handed healthy participants (range 19-65 years old, average 26 ± 11, 9 identified as female, 14 identified as male) from our university. All participants signed a consent form prior to their participation. The work was conducted in accordance with protocols approved by the Human Subjects Ethics Sub-Committee of the City University of Hong Kong. We paid 100 HKD as compensation for their participation. All subjects self-reported as healthy with no hearing impairment. Before the experiment, we examined their color and stereo vision. We used a standard Ishihara color test on a calibrated LCD screen to diagnose red-green color deficits, and a Random Dot stereo test (Stereo Optical Company Inc., Chicago, USA) set to 200 arcsec minimum level of stereopsis. We excluded three of the twenty-four participants due to color-blindness or insufficient stereo vision.

### 2.2 Augmented Reality System

To experience AR, the participants wore a Head-Mounted Display (HMD) that allowed them to see 3D computer-generated visual stimuli superimposed on their real-world view. We chose to use a Video See-Through Head-Mounted Display (VST-HMD). This device uses a pair of frontal cameras mounted on the HMD to capture the real world that is electronically blended with computer-generated imagery (Rolland et al., 1995). This allows for the creation of visually coherent virtual graphics (Collins et al., 2017; Itoh et al., 2021). We used the Varjo HMD XR-3 (Varjo, Oslo, Finland) with a pixel density of 3000 pixels per inch, a refresh rate of 90Hz, horizontal field of view 115° and vertical 80°. The dual front-facing cameras of the XR-3 have a resolution of 12 MP. We used the Valve Lighthouse tracking system to track the pose of the headset in space. To deliver sound, we used an open over-the-ear headphones (Beyerdynamic GmbH & Co. KG, Heilbronn, Germany).

Our interactive AR platform captures the volumetric representation and movement of the participant’s hand in real-time. We achieved that with the Ultraleap hand tracking sensor (Leap Motion, Inc., San Francisco, California, United States) of the Varjo XR-3 together with the Ultraleap SDK. For both conditions, as seen in Figure 1 (c) and (d), we developed a real-time fluid simulation based on the Navier–Stokes equations. For the fire condition, we created a physically correct fire simulation. The control condition differs in color as well as the fluid dynamics in a way that it would not resemble fire. We developed the whole AR experience inside the Unreal engine (Epic Games, Cary, North Carolina, USA) and used FMOD (Firelight Technologies Pty Ltd, Melbourne, Australia) for playing back realistic spatial sound effects.

### 2.3 Thermal Stimulation

To measure detection or pain thresholds, we followed the Quantitative Sensory Testing (QST) protocol of the German Research Network on Neuropathic Pain (Rolke et al., 2006). Noxious and non-noxious thermal stimuli were delivered using a commercially available thermal stimulator (Medoc TSA-2001; Ramat Yishai, Israel). This device uses a Peltier thermode with a stimulating area of 30*mm*^2^. The thermode was placed on the C7 hand dermatome (palmar of the second metacarpal) of the dominant right hand. Thresholds were determined via a method of limits. We timed the measurements in a way that they coincide with the appearance of the visuals in AR. We first measured the CDT and WDT and then the CPT and HPT. The thermal stimulator provided ramped stimuli (1°C/s) for all threshold measurements. Temperatures returned to the baseline by 1°C/s during detection threshold measurements and 5°C/s during pain threshold measurements. The next trial started after an adaptation phase of 10s at the baseline temperature. We instructed participants to indicate by pushing a button on a computer mouse with their left hand as soon as they either detected a change in the thermal sensation or until the change in temperature was perceived as painful. Cut-off temperatures were 0°C and 50°C for safety to prevent any tissue damage. Three successive trials were averaged for the detection and pain thresholds.

### 2.4 Subjective Ratings

We additionally assessed their level of motion sickness after the experiment through a Simulator Sickness Questionnaire (SSQ) (Kennedy et al., 1993). The SSQ is a self-reported list of 16 symptoms that participants are asked to rate in severity in a 4-point scale (from “absent” to “strong”). The questionnaire yields a total score and subscores on nausea, oculomotor symptoms and disorientation. It is an established method to assess symptoms after simulator use and is widely applied in virtual reality research (Davis et al., 1999; Moss and Muth, 2011; Green, 2004; Cobb et al., 1999).

A final post-study questionnaire assessed the level of presence of the experience delivered by the AR system. The goal is to assess whether they had the impression that the virtual sensory stimuli were part of the real environment. To measure the level of presence and immersion, we have adapted the presence questionnaire of (Regenbrecht and Schubert, 2021). Our presence questionnaire consists of five 7-point Likert scale questions namely *Q1*: How natural was the appearance of the fire on your right hand?. *Q2*: How much did the visual display quality interfere or distract you from observing and interacting with the visuals covering your right hand? *Q3*: Did you have the impression of seeing the virtual objects as merely flat images or as three-dimensional objects? *Q4*: Did you pay attention at all to the difference between real and virtual objects? *Q5*: Did the virtual objects appear to be on a screen, or did you have the impression that they were located on your hand)?

### 2.5 Procedures

We conducted the experiment in a sound-proof air-conditioned room, in order to keep distractions minimized. A single experimenter performed all the tests. Participants sat on a chair in front of a black table as seen in Figure 1 (a). The room temperature was maintained at 23°C. To get acquainted with the procedures, each subject performed a testing session consisting of one HPT and one CDT measurement. We presented three different stimuli in AR to all participants (See Figure 1 (b-c)), with either no additional graphics applied for baseline measurements (Figure 1 A), realistic virtual flames covering the right hand (Figure 1 B), or blue flame-like visuals covering the hand (Figure 1 C). We asked the participant to keep looking at their hand in each condition. The order of the stimuli was randomized to minimize carry-over effects (Greffrath et al., 2007; Yarnitsky and Ochoa, 1990). In each condition we measured the CDT, CPT, WDT, and HPT. We used the same order of measurements for all participants. We performed the statistical analysis in Python using the SciPy package.

## 3 RESULTS

### 3.1 Thermal Pain and Detection Thresholds

Figure 2 depicts the outcomes of the thermal QST (HPT, CPT, WDT, CDT) for all three conditions. Threshold measurements (in °C) were averaged across subjects for each of the three conditions. Because of the small sample size, determining the distribution of pain thresholds (HPT, CPT) was important for choosing an appropriate statistical method. Therefore, we performed a Shapiro-Wilk test, which showed that the CPTs could be considered normally distributed (*p >* 0.05) in all conditions. We conducted one-way repeated-measures ANOVA (one factor: *“Condition”* with three levels) on mean CPTs. Post hoc analysis was conducted with Tukey HSD tests. Since sphericity is violated for CPT (*ϵ* = 0.761), Huyn-Feldt corrected results are reported. The one-way repeated measures ANOVA showed an effect of the factor *Condition* for the CPT (*F*_1.44,28.79_, *p* = 0.007). Post hoc tests have shown that the condition *Fire* significantly increased the CPT (*p* = 0.043) (mean increase = 1.84 *±* 2.37) compared to the *Baseline* condition and compared to the *Vontrol* condition (*p* = 0.002). However, HPTs significantly diverted from the normal distribution (*p* = 0.043) during the *fire* condition and were therefore analyzed using non-parametric Wilcoxon Signed-Rank tests. They revealed that experiencing virtual flames on one’s hand in the condition *Fire* lead to a significant lower HPT (*p <* 0.001) (mean decrease: 1.55 *±* 1.66) compared to the baseline and to the control condition (*p* = 0.007). There was no significant difference in the *Baseline* and *Control* condition (*p* = 0.812).

**Figure 2.**
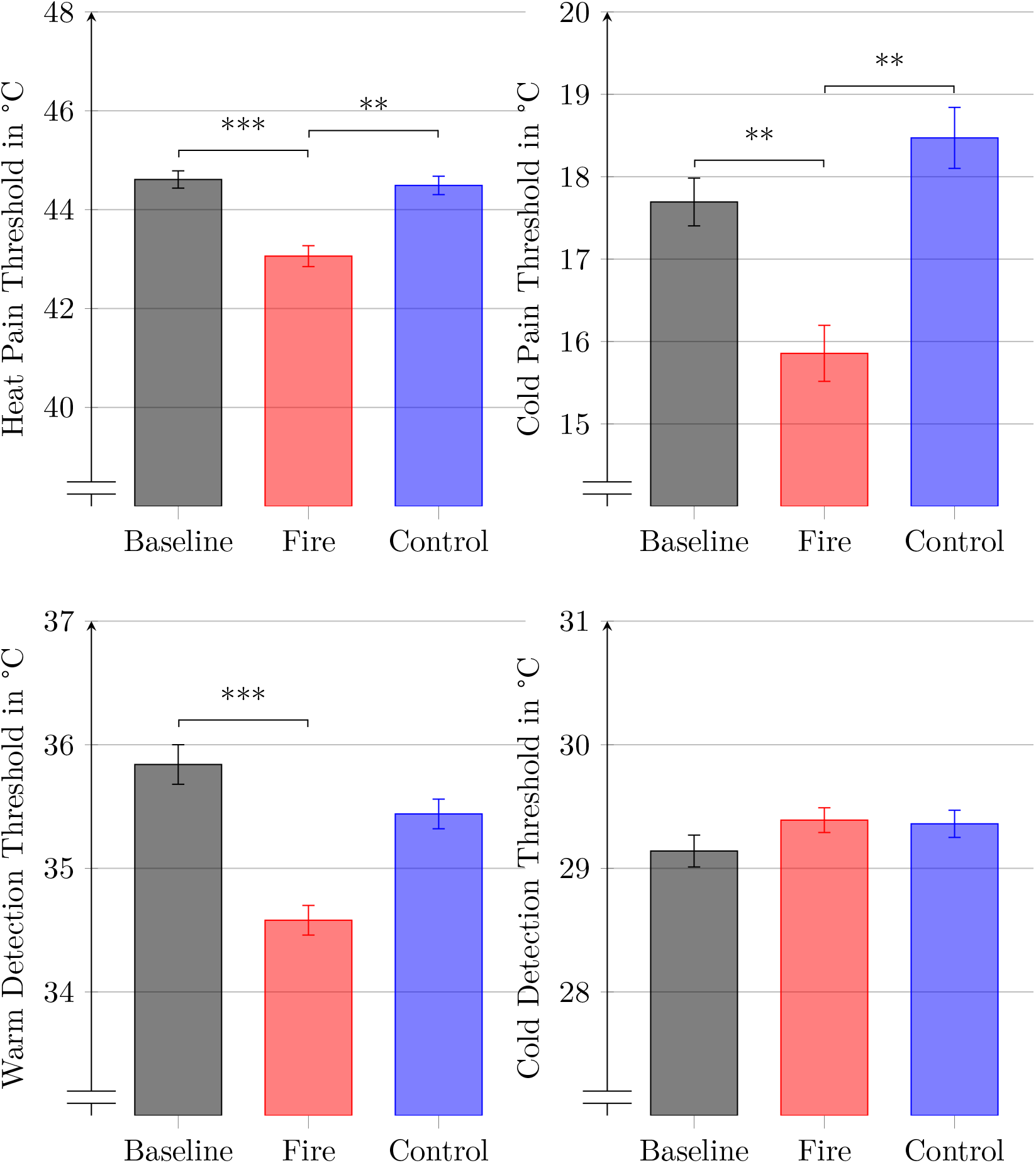
Mean thermal pain and detection thresholds for each condition. The whiskers denote the standard error. Connected bars represent significant difference (^*∗*^= *p <* .05, ^*∗∗*^ = *p <* .01, ^*∗∗∗*^ = *p <* .001).

We found the detection thresholds (WDT, CDT) to be not normally distributed (Shapiro-Wilk test, (*p <* 0.05)). In particular, WDTs were found to be positively skewed, whereas CDTs were found to be negatively skewed. These were therefore pairwise analyzed using a non-parametric Wilcoxon Signed-Rank test. The burning hand significantly decreased the WDT (*p <* 0.001). We found no other significant differences.

### 3.2 Simulator Sickness Questionnaire

The mean total score of the SSQ of 17.28 ± 3.81 indicated only little to no motion sickness effects induced by our experimental platform. *Fatigue* and *salivation increasing* were the most frequently reported symptom (*n* = 2). *General discomfort, eyestrain, difficulty focusing, nausea, fullness of the head* and *dizziness with eyes open* was reported by one participant in each case. All of the symptoms were rated as slight and none as moderate or strong.

### 3.3 Presence

On a questionnaire measuring presence in AR (7 point Likert scale), adapted from the work by Regenbrecht and Schubert (Regenbrecht and Schubert, 2021), participants reported experiencing a high degree of presence (5.30 *±* 1.40), indicating that our AR system created an effective illusion through virtual graphics that were well registered with the real world.

To explore whether the presence value and how natural the participants perceived the graphics, we calculated correlations (Pearson’s *r*) with all differences of threshold measurements compared to the baseline. However, we found no significant correlations.

### 3.4 Paradoxical Heat Illusion

Although we did not specifically ask about paradoxical heat sensations, four of the 21 participants reported the sensation of paradoxical heat. They confused a decreasing temperature ramp during a threshold measurement with an increasing one in the *Fire* condition.

## 4. DISCUSSION

The major finding of this study is that visual-auditory AR can effectively modulate nociception and thermoception of real thermal stimuli. Exposure to the burning hand in AR leads to **(1.)** a significantly lower HPT and CPT, and **(2.)** to a significantly lower WDT. The control condition does not change thresholds significantly compared to the baseline.

Our observations suggest that this AR experience can exert both hyperalgesic and analgesic effects. Seeing one’s hand on fire changes the observer’s sensitivity such that less heat is needed to produce sensations of heat pain, but also such that more cold is required to induce sensations of cold pain. In addition, less heat is needed to trigger sensation of warmth. The cross-modal effects thus modulated cutaneous perception in multiple ways. Our finding of hyperalgesic and analgesic changes is in accordance with previous studies done in VR (Martini, 2016). To the best of our knowledge, this is the first study to show a top-down modulation of HPT, CPT and WDT in AR.

There has been a big evidence for the red–warm, blue–cold association. It has been shown that children as young as about nine years of age associate the color red with warmth and blue with cold (Morgan et al., 1975). These colour-temperature correspondences can even lead to some thermoregulatory responses. Rugierri and Petruzziello found that body temperature increased significantly when participants looked through red, orange, and yellow glasses (Ruggieri and Petruzziello, 1988). Blue, green or violet did not lead to a significant change in body temperature. Moseley and Arntz have shown that a noxious stimulus, when associated with a red cue, hurts more and is actually perceived as hotter, whereas the opposite is true when associated with a blue cue (Moseley and Arntz, 2007). In a study, more closely related to our’s, by (Ho et al., 2014), the WDT was found to be lowered when the color of participants’ real hand was changed to red by a projector. A blue hand increased WDT. We were able to reproduce the results in our fire condition. However, in our case, the control condition did not significantly change any thermal detection or thermal pain threshold, even though the cue was blue. This could indicate that the dynamics of the flames lead to some perceptual ambiguity.

Our work raises the question whether, or how strongly, the location of the virtual fire affects thermal perception. Would a virtual fire at some distance from the thermal stimulator, or even just the presence of flames in the nearby environment, also lead to a change in HPTs? While (Martini et al., 2013) used a VR system that superimposes a red, blue, or green color on a virtual embodied arm to show that the reddened arm significantly decreased HPT and a blue one increased it, they also included a condition involving a virtual stimulus presented outside the limb (a coloured spot), and observed that this did not lead to a change in HPT. Their study therefore shows the importance of a virtual stimulus that is co-located with the real arm (an “embodied” experience).

Furthermore, the simple yet effective virtual stimulus employed by Martini et al. challenges the weight that realism may play in the effectivness of the modulation of thermoception. However, note that we could induce changes in pain thresholds much larger than those reported in the works of (Martini et al., 2013). Their HPT decreased by around 0.4 °C, while in our study the average decrease was 1.55 °C. We therefore hypothesize that the level of realism and intensity of the visual-auditory illusion, that is suggestive of heat, may positively correlate with how strongly cutaneous heat sensations are modulated. It was already shown that the realism and fidelity of graphics inside a virtual environment positively correlates with the level of stress response of participants (Slater et al., 2009; Weiß et al., 2021). A vivid image of flames may be particularly effective as this is universally recognized as a potential threat (Erlich et al., 2013). Additionally, this experience has been shown to elicit involuntary heat illusions in participants (Weir et al., 2013; Eckhoff et al., 2020), which might be reason for the strong modulatory effects on HPT, CPT and WDT.

The fact that we observed much greater changes in thresholds than (Martini et al., 2013) could also be due to our use of AR rather than VR. Indeed, VR requires that the participant “embodies” the virtual arm, which they may do to a greater or a lesser extent given that the VR arm is not a highly realistic representation of the participant’s actual limb. In an AR setup, the participant is able to see the real hand. (Perani et al., 2001) demonstrated that different neural networks are activated when participants observe either a real hand or a virtual hand in different degrees of realism. In the cases of the virtual hands, they found only limited differences in activation due to the different degrees of realism, compared to the response of viewing the real hand.

Our perception is shaped by an interplay of innate factors and information gained from life experience (Ernst and Bülthoff, 2004; Shams and Beierholm, 2010; Welch and Warren, 1980). We evolved in a way to respond in a specific way to specific perceptions. For example, the sound of a crackling fire is easily recognizable and well-understood, even by infants. (Erlich et al., 2013) found that infants demonstrated significantly enhanced heart rate deceleration, larger eye-blinks, and more visual orienting when listening to evolutionary fear-relevant sounds, including crackling fire. Here, seeing and listening the fire could have triggered similar reactions and thus increased the alertness and perception of pain. Our control condition had very similar visuals (brightness, fluid dynamics) and sound, yet did not modulate pain and thermal perception, thus adding further evidence that any change in pain modulation did not come from distraction or attention, rather from observing the red fire.

Interestingly, four of our participants spontaneously reported a paradoxical heat illusion during a cold stimulus so strong as to prompt them to interrupt the experiment and question the correct functioning of the thermal stimulator. This only happened in the fire condition, suggesting that the paradoxical effect is correlated to the visualization of the moving flames. Paradoxical heat illusions have been reported in as much as 35% of participants during a cold stimulus threshold experiments with pre-heating of the hand, and only 9.8% without pre-heating (Hämäläinen et al., 1982). A 19% (4 out of 21) occurrence of the paradoxical effect in our experiment may then suggest that the virtual imagery could also induce the illusion of real pre-heating, an interesting question left for further studies.

In our study, we did not observe a change in CDT. This could be since perception of cold and warm involve different neurobiological mechanisms (Patapoutian et al., 2003), presumably responding differently to cross-modal stimulation. A larger experimental sample size may be required to detect more subtle effects on the modulation of the CDT.

This study provides valuable insight into how AR can bias thermal perception. This raises some fascinating new possibilities of using AR to modulate clinical pain. AR pain treatments would be entirely non-invasive and very safe, and our results give reason to believe that they could also be quite effective.

## AUTHOR CONTRIBUTIONS

D.E. conceived the study, implemented the experimental setup and collected and analyzed the data. D.E, A.C, G.L.Y.C., and J.S interpreted the data. D.E, A.C, C.S, and J.S. designed the experiment. D.E. wrote the manuscript with the help of all authors. All authors revised the manuscript up to the final version.

